# A Dual-Phase Strategy to Integrate the VITEK MS RUO Database into Routine Clinical Practice: From Validation of Analytical Performance to Implementation of an Automated Workflow

**DOI:** 10.64898/2026.02.24.707682

**Authors:** Hua Nian, Fushun Li, Xingqi Wang, Xiaoou Yu, Jinna Dai, Yunzuo Chu

## Abstract

**Background:** Matrix-assisted laser desorption/ionization time-of-flight mass spectrometry (MALDI-TOF MS) is pivotal in clinical microbiology. The VITEK MS Research Use Only (RUO) database offers broader species coverage, yet its clinical adoption is hindered by insufficient performance validation against the approved *in vitro* diagnostic (IVD) database and inefficient manual operational workflows.

**Objective:** This two-phase study first aimed to develop and evaluate an automated integrated workflow to enhance laboratory efficiency and diagnostic capability.

**Methods:** The RUO database’s two-tier identification architecture was utilized and in-house automated relay software was developed to parse IVD results and fully automate RUO reanalysis. Phase 1 (Mar 2021–Jun 2022) involved parallel manual testing of 2,432 isolates with both databases to analyze concordance and supplementary performance. Phase 2 (Jul–Nov 2022) prospectively incorporated 3,954 isolates to implement and assess the “IVD screening – automated RUO reanalysis” workflow.

**Results:** Phase 1 demonstrated high RUO-IVD concordance (95.7% species/genus agreement). The RUO database correctly identified 98.9% of isolates and provided valid supplementary identification for 84.4% (108/128) of IVD-failed cases, with Tier 2 contributing 28.9%. Phase 2 revealed that the integrated workflow increased the overall identification rate from 95.5% to 98.7%, with Tier 2 contributing an additional 14.5%. The automated software reduced reanalysis turnaround time by > 75%, saving consumables and labor.

**Conclusion:** The VITEK MS RUO database is a reliable and complementary tool to the IVD database. Integration with automated software creates an efficient, compliant clinical workflow, providing a practical model to enhance pathogen identification for infectious disease management.

**IMPORTANCE:** The present study bridges the validation-to-application gap for the VITEK MS RUO database. We confirm its high concordance (95.7%) and complementary value to the IVD database, quantify the impact of manual workflow inefficiency and introduce an automated software solution. The strategy highlights the key role of Reference Spectra (Tier 2) in expanding coverage and simultaneously improves diagnostic efficacy (∼99% ID rate), operational efficiency (> 75% time saved) and cost-effectiveness, offering a practical model to accelerate pathogen reporting and to guide therapy.

## Introduction

Matrix-Assisted Laser Desorption/Ionization Time-of-Flight Mass Spectrometry (MALDI-TOF MS) has become a cornerstone technology for pathogen identification in modern clinical microbiology laboratories. Its core strengths lie in its ability to provide rapid, high-throughput, low-cost and accurate species-level identification, fundamentally transforming the landscape of microbiological diagnosis (1–3). Currently, commercial MALDI-TOF MS platforms, represented by systems such as VITEK MS, are widely used globally. Their integrated and rigorously validated *in vitro* diagnostic (IVD) databases, approved by regulatory agencies (e.g., the U.S. FDA), serve as the authoritative basis and gold standard for issuing clinical reports (4–6). However, with the increasing complexity of the clinical infectious disease spectrum – particularly in the context of immunocompromised patients, complex surgeries and the widespread use of broad-spectrum antibiotics – the isolation rates of anaerobic bacteria, Gram-positive bacilli, rare yeasts and atypical pathogens continue to rise (7–9). These microorganisms are often not covered or incompletely covered by standard IVD databases, creating a significant “blind spot” in current rapid clinical microbiological identification (10, 11).

To address this challenge, instrument manufacturers also provide Research Use Only (RUO) databases, which offer broader microbial coverage and more frequent updates (11). Theoretically, these databases are ideal tools for expanding laboratory identification capabilities and tackling challenging pathogens. However, their clinical application has long faced two major bottlenecks, forming a translational gap between “research potential” and “routine use.” First, there is a lack of evidence and compliance concerns. Although RUO databases have superior species coverage, there is a lack of prospective, head-to-head comparative performance data between RUO and the approved IVD databases using the same clinical strains. Are the identification results for overlapping lineages sufficiently consistent? What is the actual accuracy rate of supplementary identifications? Without clarity on these critical performance parameters, laboratories lack the scientific basis and confidence to use RUO results to support clinical decision-making, restricting their use strictly to research settings. Second, there is an efficiency bottleneck in the workflow. Under current operational protocols, if an RUO database is used to retest strains that failed identification using the IVD database, it requires technicians to perform cumbersome manual steps: screening failed samples, recording target plate positions, manually switching modes in the software and recreating detection templates. This process is not only time-consuming (often taking tens of minutes) and error prone but also extremely difficult to implement routinely in high-volume laboratories. As a result, this powerful tool is often sidelined and fails to realize its full clinical potential.

Therefore, to truly unlock the diagnostic potential of RUO databases, these two issues must be addressed systematically. This study aims to construct a comprehensive solution through a two-phase, sequential prospective investigation. In the first phase, we will perform synchronous parallel testing using both IVD and RUO databases on a large cohort of clinically isolated strains. This phase seeks to answer the core performance questions: What is the level of agreement between the two databases? How effective and accurate is the RUO database in supplementing identification where the IVD database falls short? In the second phase, based on the performance validation results from the first phase, we will independently develop an automated switching software module. This will be followed by a prospective evaluation of the integrated workflow – “IVD primary screening → automated RUO supplementation” – in a real-world clinical setting. The central objective is to verify whether this workflow can achieve high efficiency and routine applicability while ensuring the reliability of results, thereby substantially enhancing the overall diagnostic capability of the laboratory.

## Materials and Methods

### Study Design and Isolates

Bacterial isolates were obtained from various culture types collected for routine patient care at the First Hospital of China Medical University. These included anaerobes, fastidious organisms, non-fermenting bacteria, Enterobacteriaceae, Gram-positive bacilli, Enterococcus spp., Staphylococcus spp., and Saccharomyces spp. Specimen types encompassed Bone marrow, tissue, sterile body fluids, wounds, abscesses, respiratory samples and blood cultures. This prospective two-phase study was conducted at this large academic medical center (**Fig. 1**). Phase 1 (March 2021 to June 2022) involved parallel head-to-head testing of 2,432 non-duplicate clinical isolates using the FDA-approved VITEK MS IVD database (v3.2) and the RUO database (v4.16), primarily to evaluate performance concordance and supplementary identification capability. Phase 2 (July to November 2022) prospectively enrolled 3,954 consecutive isolates to validate an automated “IVD primary screening – RUO supplementation” workflow enabled by in-house relay software (ver. 3.1; **Fig. 2A**). Isolates were collected from key clinical departments (e.g., Surgical ICU, Organ Transplant) and diverse specimen types (e.g., sputum, urine, blood, drainage fluid).

**FIG 1.**
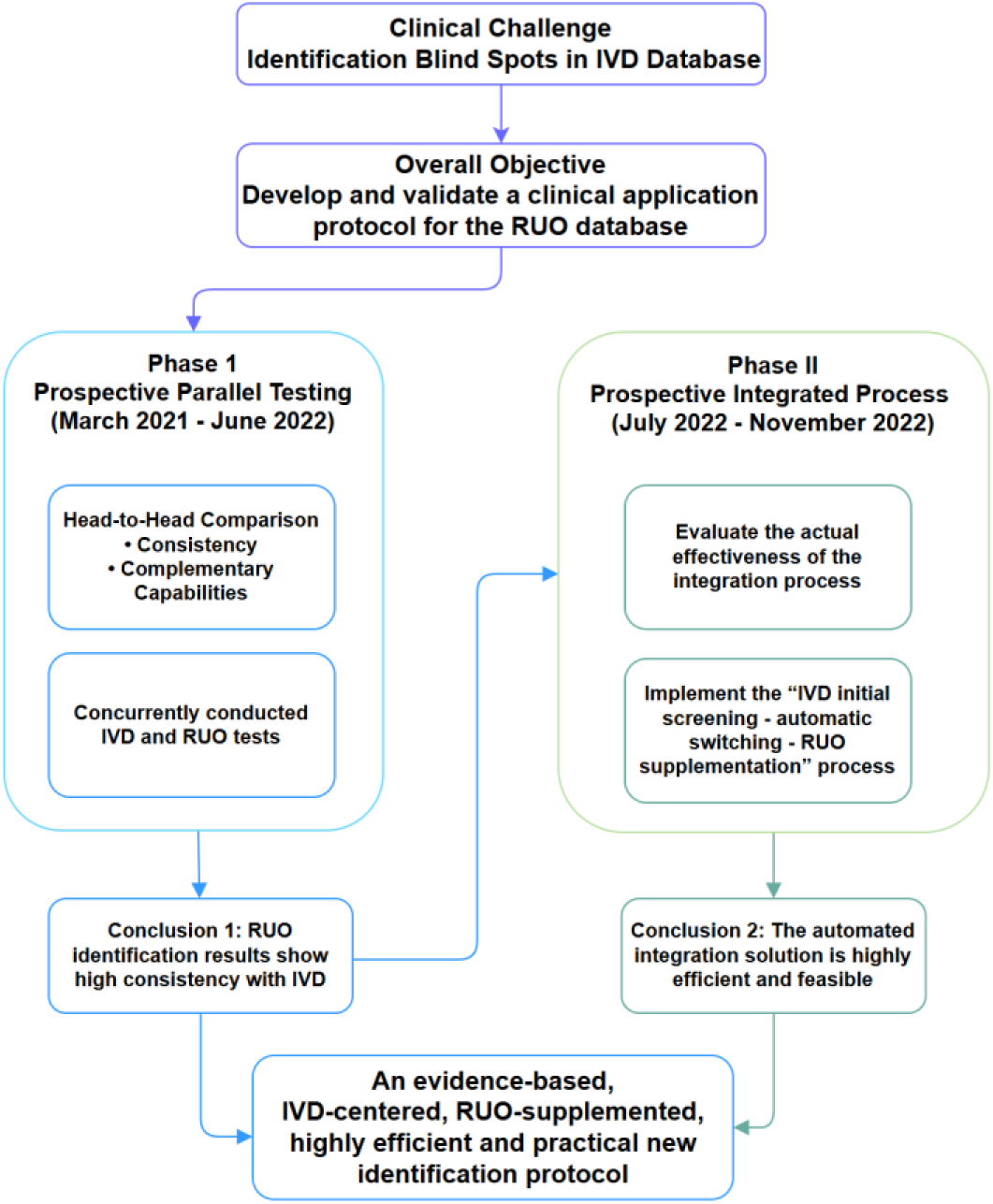
Schematic flowchart of the consecutive two-phase prospective study design. IVD, *in vitro* diagnostic; RUO, Research Use Only.

**FIG 2.**
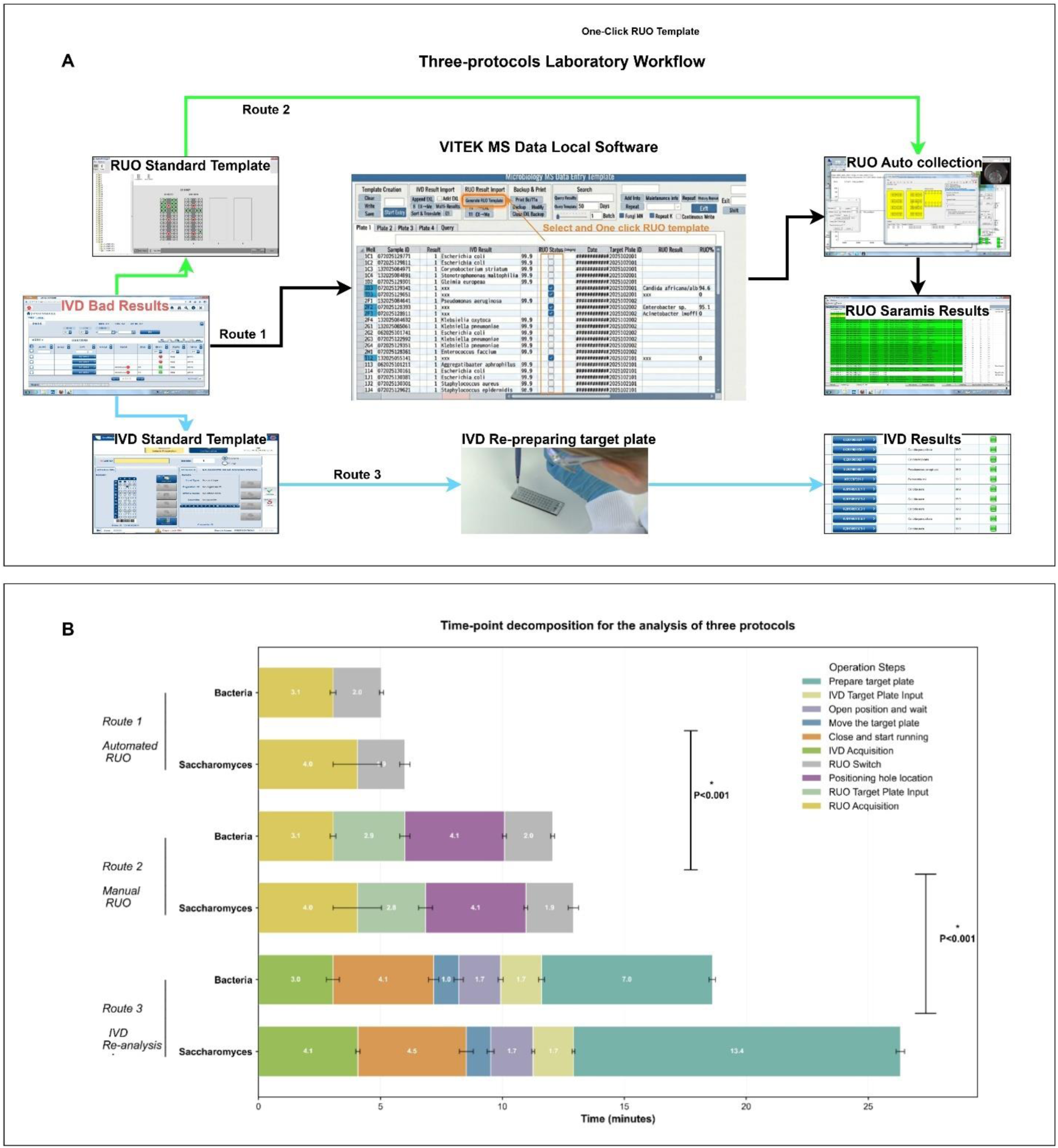
Detailed workflow of the three re-analysis protocols. **(A)** Schematic overview of the three protocols: Route 1 (Automated RUO), Route 2 (Manual RUO), and Route 3 (Regular IVD re-analysis). **(B)** Time-point decomposition for the analysis of five sample spots and one quality control spot across all protocols. Ket record steps include: Prepare target plate, IVD target input, Open position and wait, Move the target plate, Close and start running, IVD acquisition, RUO switch, Positioning hole location, RUO target plate input, and RUO acquisition. IVD, *in vitro* diagnostic; RUO, Research Use Only. Error bars represent the standard deviation.

### MALDI-TOF MS Analysis

Isolates, including anaerobes and fastidious organisms, were cultured using standard media and conditions (12). For MS analysis, fresh colonies were spotted onto a VITEK MS target plate. Bacterial spots were overlaid directly with 1 µL CHCA matrix, while yeast spots first received 1 µL of 70% formic acid for on-target extraction. Spectral acquisition and analysis were performed on a calibrated VITEK MS system with daily quality control using *Escherichia coli* ATCC 8739, *Klebsiella aerogenes* ATCC 13048 and *Candida glabrata* ATCC MYA-2950.

### Databases and Identification Criteria

Identifications from the IVD database (1,316 species) were reported via MYLA software; a confidence score > 60% defined a species-level identification. The RUO database (2,034 species, analyzed via SARAMIS v4.16) employs a two-tier hierarchical system. Tier 1 (SuperSpectra) provides direct species identification with a similarity score ≥ 75%. If this threshold is not met, the system proceeds to Tier 2 (Reference Spectra), where an identification is accepted with a score > 50% (or > 40% for consecutive same-species isolates from one source) when corroborated by phenotypic data.

For isolates that failed identification using the IVD database, a comprehensive review and verification process was initiated. This included re-examination of the original colony morphology, assessment for single-colony purity. If necessary, isolates were subcultured to obtain pure growth and then re-analyzed using the IVD database(13). Discrepancies between databases or identification of rare species by the RUO database were resolved through molecular sequencing. Bacterial and fungal isolates were analyzed via 16S rRNA and 28S gene sequencing, respectively. The raw sequence (fasta) data were deposited in the National Center for Biotechnology Information (NCBI) under accession number (Table S1,Table 3). Sequences were compared using NCBI BLAST, with the highest similarity and reliable annotation accepted as definitive.

### Implementation Workflow

In phase 1, each isolate underwent sequential analysis: initial IVD identification followed by manual instrument switching and RUO re-analysis of the same spot. In phase 2, all isolates underwent routine IVD screening. For those failing IVD identification, the in-house automated relay software parsed the result file, identified target coordinates of failed samples, regenerated a RUO-specific test template and switched the instrument mode to perform RUO re-analysis automatically.

### Efficiency

To quantitatively evaluate operational efficiency, we analyzed and compared three distinct specimen re-analysis pathways (**Fig. 2A**). A representative panel of test strains that two bacteria (*Staphylococcus aureus* as spot 1 and *Escherichia coli* as spots 2–5), one yeast ( *Candida albicans* as spot 1-5), and the quality control strain E. coli ATCC 8739 was selected for a timed analysis. The precise time points for key procedural steps were recorded for each pathway (**Fig. 2B**). The three re-analysis pathways were defined as follows: Route 1 (Automated Software-Mediated RUO Re-analysis): The in-house developed relay software automatically parsed the initial IVD results, generated a new target template for RUO analysis, and switched the VITEK MS instrument to RUO mode for data acquisition. Route 2 (Manual RUO Re-analysis): The operator manually located the target spot coordinates, created a new RUO analysis template, and then switched the instrument to RUO mode for data acquisition. Route 3 (Full Re-preparation with IVD Re-analysis): A completely new target spot was prepared from a fresh colony, and the standard IVD workflow was repeated for identification. To ensure robustness and account for operator variability, three independent technicians each performed the entire procedure for all three routes, with three technical replicates per route per operator. Time savings were calculated as: time for “Route 3” minus “Route 1” or “Route 2”, respectively. (Table 4)

### Statistical Analysis

All identification results from both the IVD and RUO were recorded for each isolate, species/genus-level accuracy and inter-database concordance rates were calculated. Differences in hands-on time between workflows were analyzed using one-way ANOVA followed by Tukey’s post hoc test, with *P* < 0.05 considered significant. Data analysis utilized Python 3.9 and GraphPad Prism 10.

## Results

### Phase 1: Prospective Parallel Testing Results

#### Strain Inclusion and Database Coverage

A total of 2,432 strains were prospectively analyzed. Both databases covered 2,397 strains (98.6%). Individually, the IVD database covered 2,400 strains (98.7%) and the RUO database covered 2,408 strains (99.0%).

#### Head-to-Head Identification Performance

For the 2,397 shared isolates, the IVD database achieved a species-level accuracy of 94.79%, a genus-level accuracy of 1.21% and a failure rate of 4.01%. Using its hierarchical system, the RUO database achieved a species-level accuracy of 95.58% (Tier 1: 91.49%; Tier 2: 4.09%) and a genus-level accuracy of 3.09%, with a lower failure rate of 1.08% and no misidentifications. Performance varied by microbial group. Anaerobes, Enterobacteriaceae, Enterococcus spp., and Staphylococcus spp. were accurately identified by both systems. For non-fermenting bacteria, species-level accuracy was similar, but RUO showed better genus-level performance and lower failure rates. Gram-positive bacilli were most challenging (10); the IVD database achieved 76.3% species-level accuracy with high failure, while RUO increased reportable results to 81.4% via Tier 1 (45.8%) and Tier 2 (35.6%) identifications. For yeasts, RUO accuracy (95.9%) was slightly higher than IVD (89.4%). For fastidious organisms, both databases achieved 95.2% accuracy, with RUO results comprising Tier 1 (85.7%) and Tier 2 (9.5%) identifications (1, 14).

The two databases showed high overall concordance for the 2,397 shared isolates, with species-level agreement of 91.7% (2,188/2,397) and taxonomically acceptable (genus-level or above) agreement of 95.7% (2,285/2,397). Discordant results accounted for 4.27%, while mutual identification failures were rare (0.42%). Concordance varied by microbial group, being highest for anaerobes (97.0%) and lowest for Gram-positive bacilli (63.2%) at species level.

#### Analysis of the Supplementary Identification Efficacy of the RUO Database for IVD-Failed Isolates

Among the 128 isolates not identified by the initial IVD procedure, the IVD database covered 96 isolates, while the RUO database covered 107. The RUO database yielded definitive identification (to species or genus level) for 110 isolates, of which 2 were incorrect. Thus, 108 correct supplementary identifications were achieved, corresponding to a supplementation rate of 84.4% (**Table 2**). Analysis attributed 26 (20.3%) of failures to initial “detection mode selection” error (**Table 2**).

**TABLE 1.**
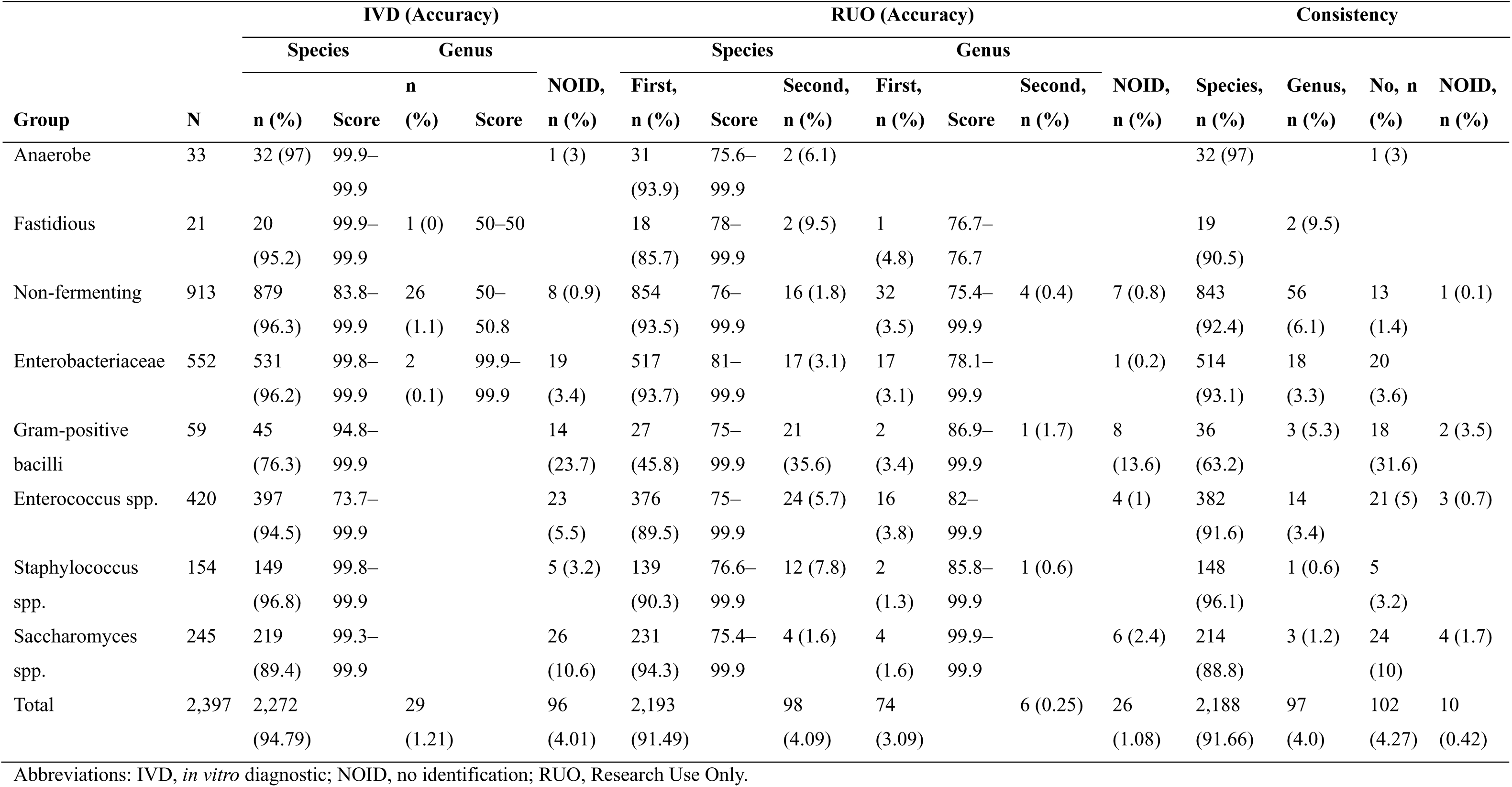
Detailed comparison of the identification performance of the IVD and RUO databases for different microbial categories (n = 2,397)

**TABLE 2.**
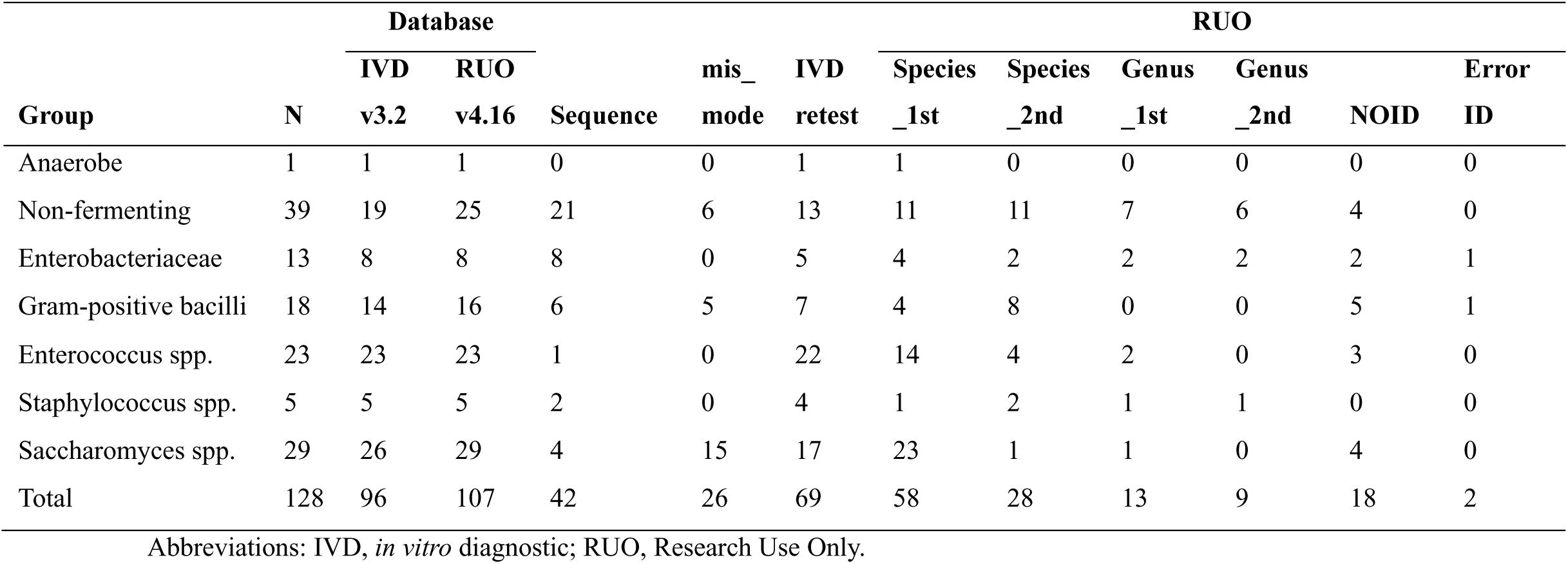
Detailed analysis of 128 isolates with failed initial IVD identification and the supplementary efficacy of the RUO database.

Of the successful supplementary identifications, 58 (45.3%) were achieved by Tier 1 (SuperSpectrum) at species level, while Tier 2 (Reference Spectrum) contributed 28 (21.9%) at species level. Overall, Tier 2 enabled reporting for 37 isolates (28.9% of all IVD failures).

Non-fermenting bacteria and yeasts constituted most failures. The RUO database successfully identified 35 (89.7%) of non-fermenters and 25 (86.2%) of yeasts, with Tier 2 playing a greater role in non-fermenter 17 (43.6%) than yeast 1 (3.4%) identification.

For 32 isolates belonging to species absent from the IVD library, the RUO database formally covered 6 species (11 isolates) and correctly identified 5 species (9 isolates). For remaining species not formally listed, presumptive genus-level identification was achieved for 15 isolates, albeit with 2 errors (**Table 3**).

**TABLE 3.**
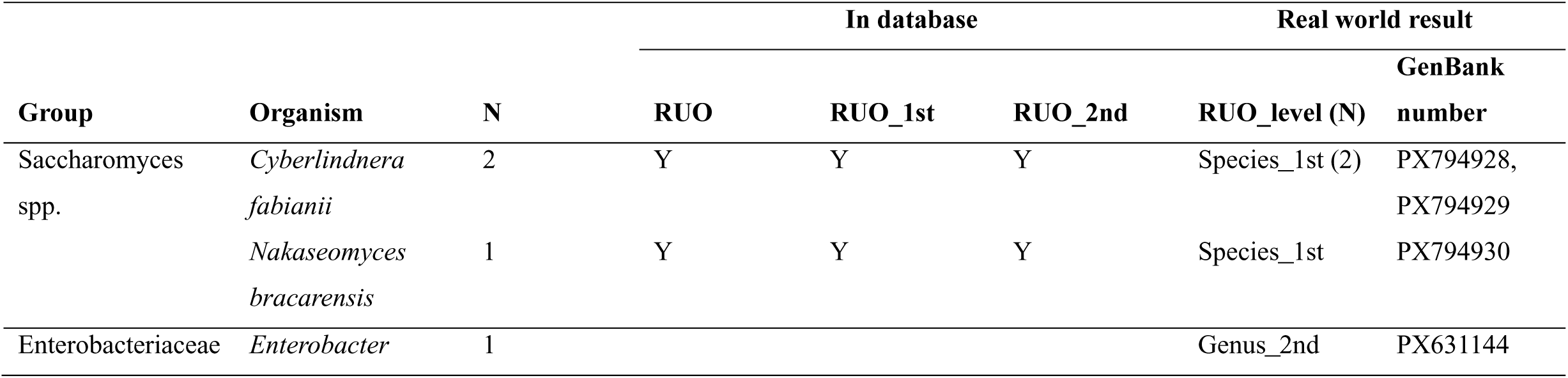
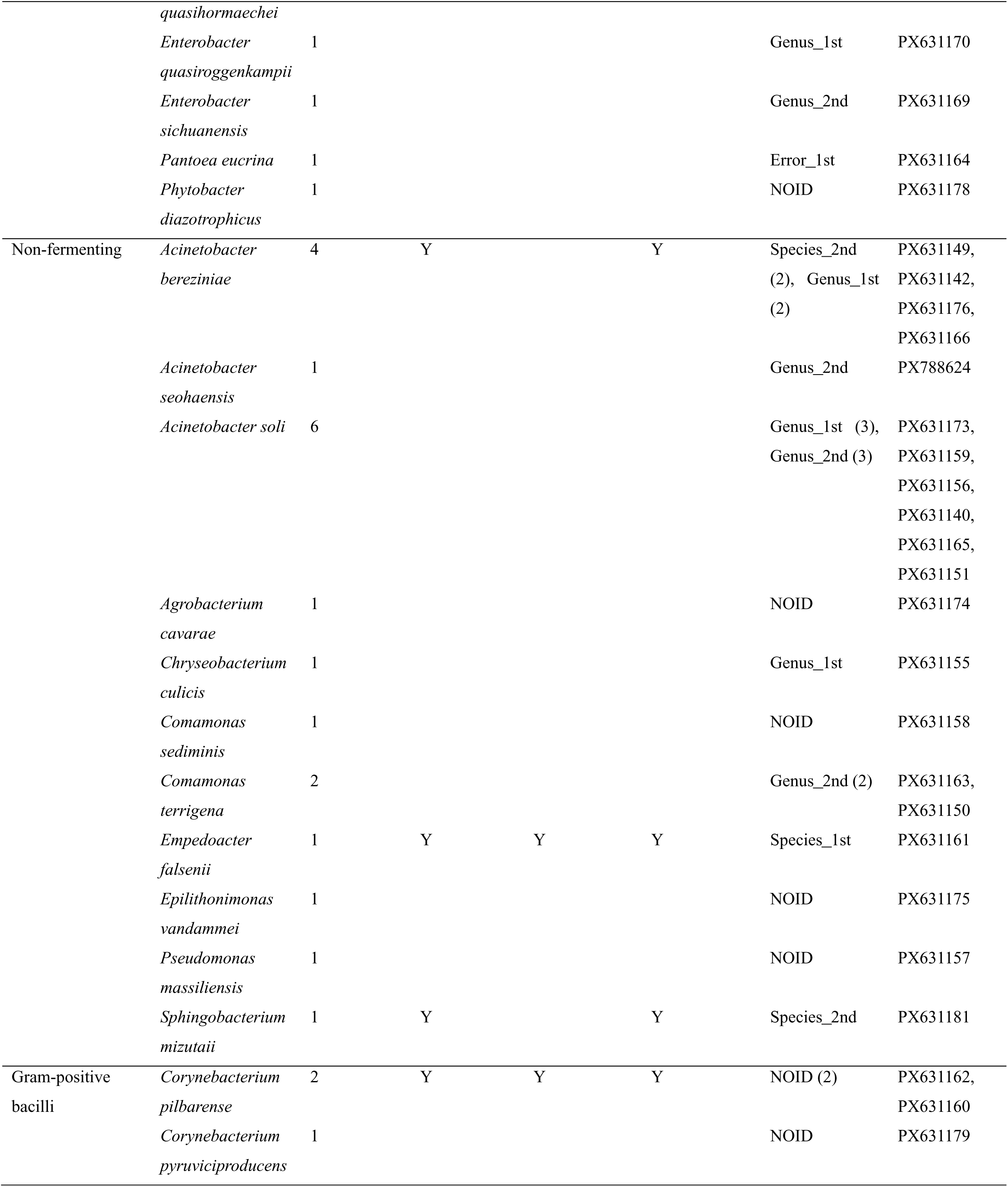
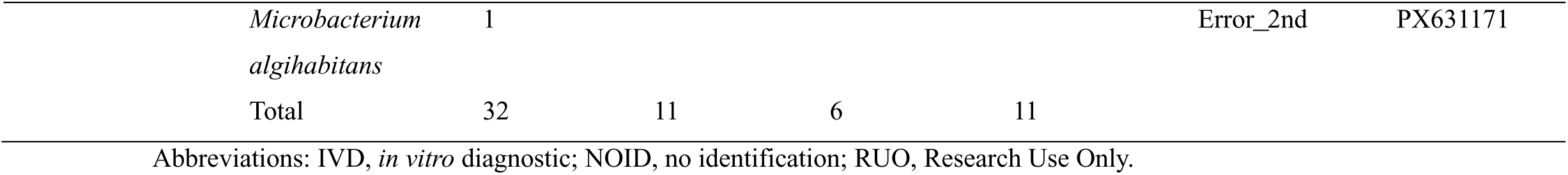
List of 32 bacterial strains absent from the IVD standard database.

#### Panoramic Performance Comparison of IVD and RUO Databases Based on a Sankey Diagram

The Sankey diagram (**Fig. 3**), constructed from the parallel testing data of 2,432 isolates, provides a comprehensive visualization of the workflow and outcomes for both databases. It shows that the IVD database identified 2,275 isolates to the species level (IVD_species), while 128 isolates yielded no identification (IVD_NOID). In contrast, the RUO database, utilizing its hierarchical system, identified 2,197 isolates to species level via Tier 1 (RUO_species_1st) and supplemented an additional 101 isolates via Tier 2 (RUO_species_2nd). Of the 128 isolates that failed IVD identification, the RUO database successfully reclassified the majority to the species or genus level. Only 34 isolates remained without a result (RUO_NOID), and 2 misidentifications were recorded. This visualization intuitively demonstrates that the RUO database, through its tiered identification strategy, maintains a high species-level identification rate while significantly reducing the rate of non-identification, thereby achieving more comprehensive microbiological coverage.

**FIG 3.**
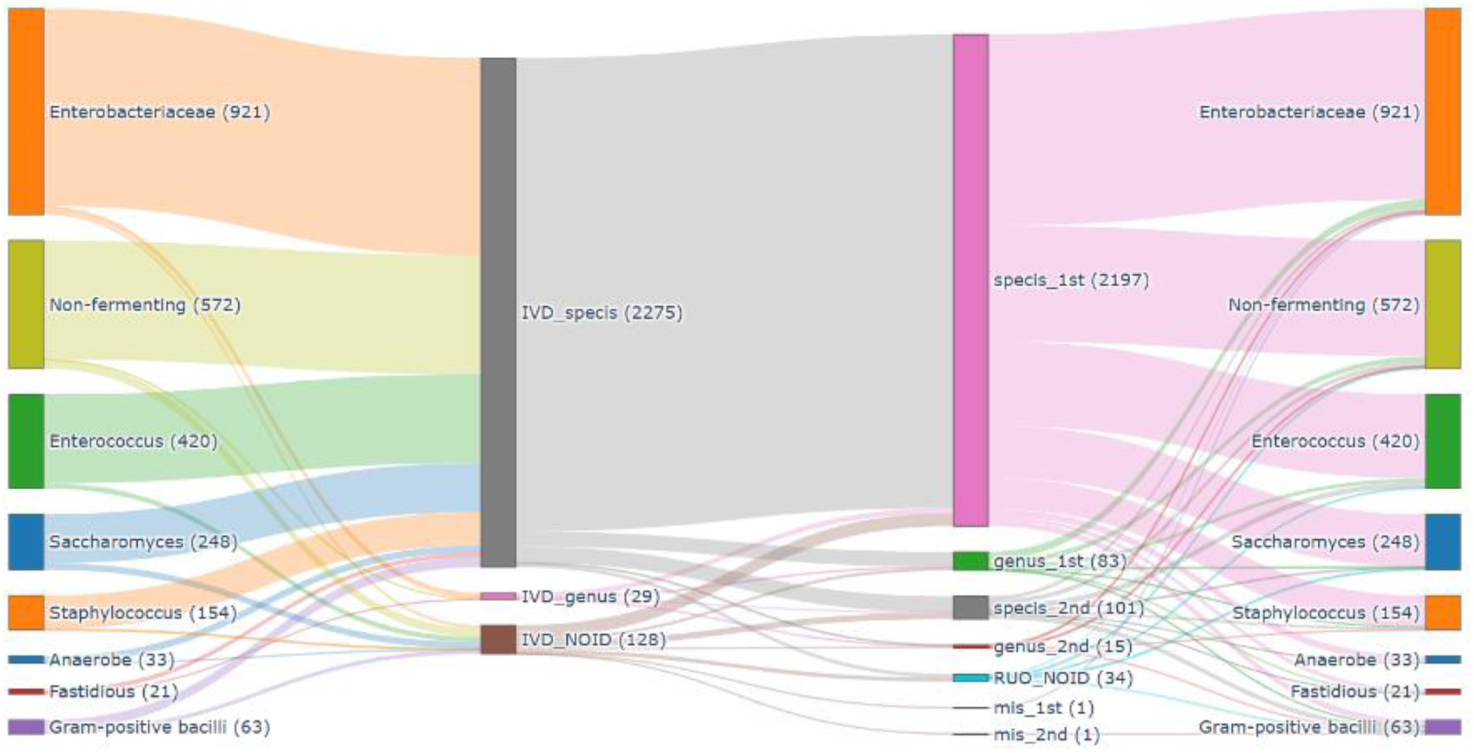
Flowchart of microbial identification results for phase I (2,432 strains). The left side displays the total bacterial strain count. After passing through the two parallel identification systems (IVD and RUO) in the middle, strains are categorized based on identification results (species, genus, inconclusive, error) and spectral level (determined by RUO identification), converging at the endpoint on the right. Line width is proportional to the corresponding strain count. IVD, *in vitro* diagnostic; NOID, no identification; RUO, Research Use Only.

#### Efficiency Comparison of the Three Re-analysis Protocols

Three re-analysis workflows were compared as outlined in **Fig. 2A**. Time-motion analysis of each step (**Fig. 2B**) quantified efficiency gains with the RUO/IVD system (**Table 4**). Overall, the total process completion time was different between Routes (Route 1 (automated software) : 5.04 min (bacteria) and 6.0 min (yeasts); Route 2 (manual) : 12.07 and 12.92; Route 3 (full re-preparation) : 18.61 and 26.33, respectively), with the Route 1 (−13.57 min (bacteria), −20.32 min (yeasts), P < 0.001, one-way ANOVA/Tukey’s) and Route 2 (−6.54 min (bacteria), −13.41 min (yeasts), P < 0.001, one-way ANOVA/Tukey’s) being faster than the Route 3. Compared to Route 3, Route 1 saved 72.9% (bacteria) and 77.2% (yeasts) in time; Route 2 saved 35.15% and 50.93%. Additionally, Route 3 also consumed substantially more resources, using 16 target spots and ∼$60 in reagents for only 5 samples.

**TABLE 4.** Workflow efficiency comparison.

| Evaluation metrics |  | Route 1: Auto RUO |  | Route 2: Manual RUO |  | Route 3: Regular IVD |
| --- | --- | --- | --- | --- | --- | --- |
|  |  | minutes±std | % | minutes±std | % |  |
| Test time | Bacteria | 5.04±0.72 |  | 12.07±1.11 |  | 18.61±0.53 |
|  | Saccharomyces spp. | 6.0±0.28 |  | 12.92±0.35 |  | 26.33±0.78 |
| Time-savings compared to Route 3 | Bacteria | 13.57 | 72.90 | 6.54 | 35.15 | - |
|  | Saccharomyces spp. | 20.32 | 77.20 | 13.41 | 50.93 | - |
| Operational complexity |  | Low |  | Medium |  | High |
| Workflow integration |  | Excellent |  | Poor |  | - |
| Batch samples |  | Simple |  | Complex |  | Complex |
| Target consumption (price) | | 0 | | 0 | | 16 (≈ \$60) |

Abbreviations: IVD, *in vitro* diagnostic; RUO, Research Use Only.

#### Phase 2: Prospective Performance Validation of the Integrated Automated Workflow

Building on the performance verification of the RUO database from phase 1, the objective of phase 2 was to prospectively assess the practical effectiveness of integrating the RUO database into the routine workflow via the in-house developed automated switching software. This phase prospectively included 3,954 consecutive clinical isolates. The results of the initial screening using the standard IVD database are summarized in **Fig. 4**. The IVD database successfully identified 3,526 isolates to the species level and 249 isolates to the genus level, yielding an initial overall identification rate of 95.5% (3,775/3,954). The remaining 179 isolates (4.5%) failed initial IVD identification. These 179 IVD-failed isolates were seamlessly subjected to supplementary analysis using the RUO database via the automated workflow. The RUO database successfully provided valid identification for 126 of these isolates, achieving a supplementary identification rate of 70.4% (126/179). Specifically, 100 isolates were identified to species level and 26 to genus level, with Tier 2 (Reference Spectrum) identification contributing 26 isolates (14.5%) (**Table S2**). This integrated strategy, which prioritizes the IVD standard, resulted in a significant increase in the laboratory’s overall identification rate from 95.5% to 98.7%, representing an absolute improvement of 3.2 percentage points.

**FIG 4.**
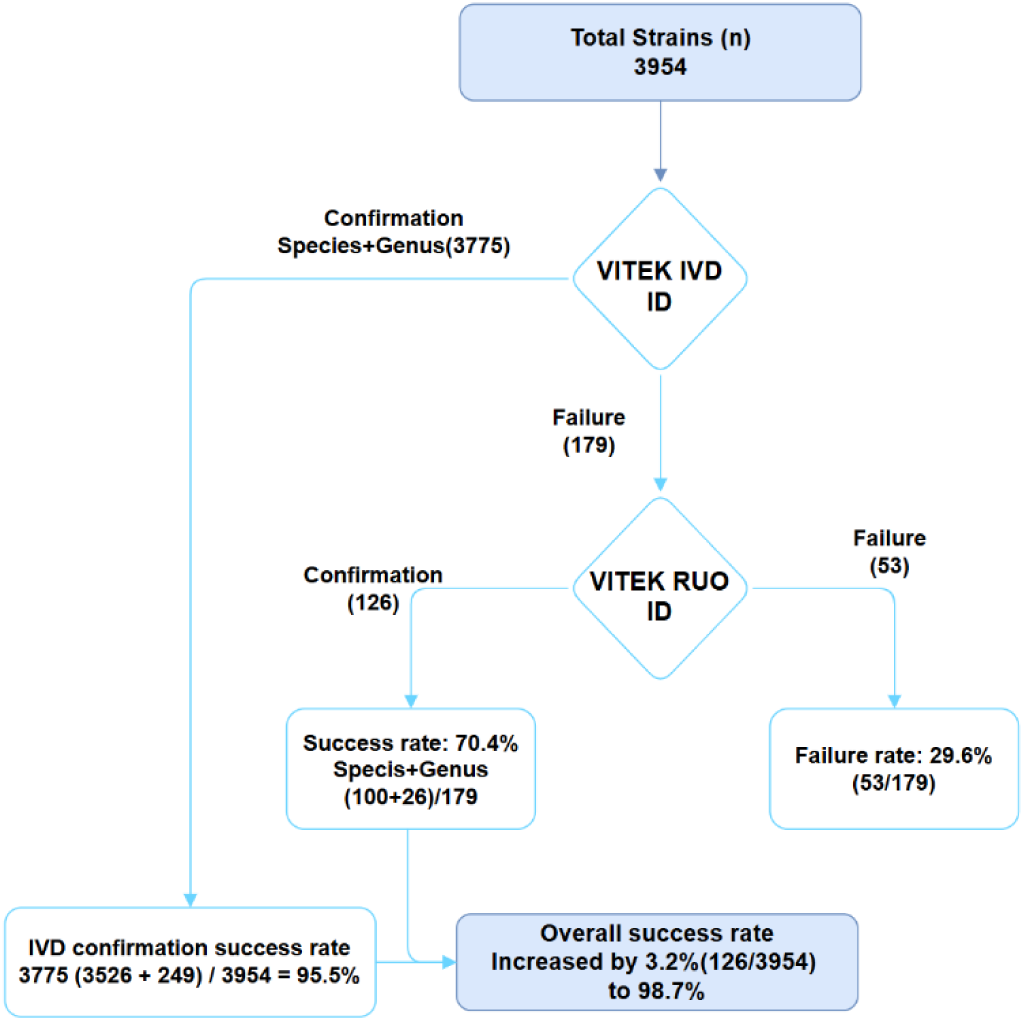
Practical performance of the integrated automated workflow in phase 2. IVD, *in vitro* diagnostic; RUO, Research Use Only.

## Discussion

This study establishes a complete evidence chain from performance validation to clinical implementation. Unlike most prior comparisons of RUO databases with standard methods (11, 13, 15, 16), we employed a two-phase prospective design. This not only confirmed high species/genus-level concordance (> 95%) and substantial supplementary identification rates (∼70–84%) between the RUO and FDA-approved IVD databases but, crucially, via in-house automated software, translated this potential into a practical routine workflow. This efficient and stable integration raised the laboratory’s overall identification rate to 98.7%, offering a methodological model to overcome the common “easy-to-validate but hard-to-implement” challenge in advanced diagnostics.

In a head-to-head comparison of 2,397 clinical isolates, this study confirmed that both the RUO and FDA-approved IVD databases exhibit high identification capability (overall correct ID rates: IVD 96%, RUO 98.9%) and show substantial equivalence, with > 91% species-level agreement. This supports the “clinical-grade” reliability of the RUO database and provides a technical basis for its use in defined scenarios.

The two databases differ in emphasis: the IVD database offers stable, “ready-to-use” identification for common pathogens (e.g., Enterobacteriaceae), suited for routine high-throughput workflows. In contrast, the RUO database, with its larger spectral library and flexible hierarchical system (especially Tier 2 spectra), provides broader coverage and lower failure rates for challenging groups such as Gram-positive bacilli.

Notably, the RUO database successfully identified 84.4% of isolates that failed initial IVD identification, spanning Enterococcus spp., non-fermenters, yeasts, and rare Gram-positive bacilli and anaerobes. This complements earlier views on MALDI-TOF MS limitations for certain groups (17) and indicates that single-run IVD success can be affected by technical variables; RUO integration effectively bridges this gap (18).

This study is the first to systematically evaluate the clinical value of secondary reference spectra (Tier 2) in the VITEK MS RUO database. Unlike prior reports that focused mainly on the primary library (SuperSpectra/Tier 1), we quantified Tier 2’s contribution to challenging identifications. In phase 1, 4.09% of all species-level identifications relied solely on Tier 2 matches, moving beyond the “single best match” paradigm to highlight the strength of its hierarchical algorithm. This contribution was particularly evident among difficult-to-identify organisms. For example, Tier 2 enabled species- or genus-level reporting for 28.9% of successfully supplemented identifications (**Table 2**). Operationally, when a spectrum did not match a SuperSpectrum with high confidence, the system compared it against a larger, more inclusive Tier 2 library containing strain variants, related species and weaker spectral profiles. Through probabilistic matching, it provides reliable genus- or species-level inferences. Our findings validate that this hierarchical design effectively expands “reportable results” by converting isolates with atypical profiles into clinically useful identifications. This confirms that a tiered identification architecture enhances both system robustness and coverage – a key insight for developing intelligent multi-tier algorithms to address microbial diversity.

This study is also the first to systematically identify and quantify the impact of the pre-analytical human error of “incorrect detection mode selection” on the success rate of IVD identification. The IVD workflow requires operators to pre-select either a “bacteria” or “fungi” testing mode. An erroneous selection, often due to atypical colony morphology, directly leads to identification failure by the IVD database.

Analysis of the 128 isolates that failed initial IVD identification (**Table 2, Table S3**) revealed specific scenarios: 15 yeasts growing on blood agar were easily misjudged as bacteria, while 11 non-fermenting bacteria and Gram-positive bacilli growing on Sabouraud or CHROMagar™ medium were sometimes mistaken for yeasts, triggering the wrong mode selection. In contrast, the RUO database employs a unified acquisition and analysis workflow, eliminating this pre-selection step and thereby preventing such human errors. The analysis showed that 26 out of the 128 IVD failures (20.3%) were attributable to “incorrect detection mode selection”. The RUO database successfully identified 25 of these 26 isolates (the single failure was a *Cryptococcus neoformans* isolate, where the use of the bacterial mode without formic acid resulted in a poor-quality spectrum). This finding not only confirms the technical advantage of the RUO workflow but also demonstrates its practical value in correcting this specific type of pre-analytical human error.

The RUO database precisely supplements specific IVD “blind spots.” As shown in **Table 3**, it successfully identified 9 isolates across 5 clinically significant species: rare pathogenic yeasts *Cyberlindnera fabianii* (19–21) and *Nakaseomyces bracarensis* (22); *Acinetobacter bereziniae* (non-*baumannii* complex) (23); and opportunistic pathogens *Empedobacter falsenii* (24) and *Sphingobacterium mizutaii* (25). This targeted supplementation enhances diagnostic capability for emerging, rare and healthcare-associated pathogens, supporting precision therapy and outbreak investigation. Among IVD-failed isolates not formally listed in the RUO database, 15 of 21 received presumptive genus-level identification based on spectral similarity, though with 2 misidentifications (**Table S4**).

The Sankey diagram (26), constructed from the parallel testing data of 2,432 isolates, provides a macroscopic, data-flow perspective for the intuitive comparison of the identification logic and performance between the IVD and RUO database (**Fig. 2**). This visualization confirms that the vast majority of isolates (> 93%) can be identified to the species level by either system, yet their pathways differ markedly. The IVD output is concentrated in two primary categories: “species Identification” (2,275 isolates) and “no result” (128 isolates), a straightforward dichotomy that facilitates clinical decision-making. In contrast, the RUO system reveals a refined hierarchical structure: alongside the 2,197 isolates directly identified via Tier 1 (SuperSpectrum), an additional 101 isolates achieved species-level identification through Tier 2 (Reference Spectrum). The chart clearly demonstrates that the 128 isolates failing IVD identification are the primary target for the RUO system’s supplementary value. The majority were successfully converted into valid species- or genus-level results, leaving only 34 isolates with no result and an extremely low misidentification rate (2 isolates). This qualitative panoramic view strongly corroborates the core finding of this study: through its hierarchical identification system, the RUO database maintains a high species-level identification rate while significantly converting IVD’s “no result” isolates into taxonomically informative data with diagnostic value, thereby achieving more comprehensive isolate coverage. Objectively, the diagram replicates the complementary relationship between the two strategies from a workflow perspective: their results show high overlap within the shared taxonomic spectrum, whereas for IVD blind spots, RUO provides effective, multi-tiered supplementation.

This time-motion analysis confirms that integrating the RUO database markedly improved workflow efficiency and reduced turnaround time. Compared with full IVD re-preparation (Route 3), the automated RUO re-analysis (Route 1) saved 75.05% of the time for bacteria and 78.85% for yeasts. This acceleration facilitates earlier reporting and enables more timely adjustment of antimicrobial therapy (27, 28), which is particularly meaningful in critical care and invasive fungal infections. Economically, Routes 1 and 2 avoid the need for additional target-plate spots. Processing five samples via Route 3 required a full 16-spot segment, whereas Routes 1/2 used no extra spots, yielding an estimated consumable saving of about $40 per run and reducing associated reagent and labor costs. Operationally, this integrated workflow also provides resilience: if IVD-mode QC fails, the system can switch directly to RUO mode, preventing repeat testing and further conserving resources.

The supplementary capability of the RUO database derives from multiple technical and operational distinctions discussed in the literature. First, its spectral library provides broader species coverage, encompassing more strain variants, rare organisms and emerging pathogens, thereby directly expanding the identifiable microbial spectrum (4, 29). Second, the matching algorithms and decision thresholds differ: the RUO system employs a hierarchical, probabilistic strategy with more inclusive thresholds, enabling species- or genus-level inference even when the IVD system reports “no identification.” Significant contributing factors also include pre-analytical and analytical variations: (1) “mode selection error” arising from manual choice of detection mode; (2) the “sweet-spot effect” on the MALDI target plate, where positional differences affect ionization efficiency and spectrum reproducibility (30); and (3) operator- and culture-related variables such as incubation time and sample preparation technique. These factors introduce spectral variability that can differentially affect matching outcomes between the two systems. In summary, the RUO database’s supplementary performance results from the interplay of its richer spectral library, more flexible algorithm design and the technical variabilities inherent in the testing process. When implemented with a standardized protocol and supported by automated software to minimize human selection bias, it serves as a stable and reliable supplemental identification tool.

This study had several limitations. First, it was conducted at a single center and the spectrum of isolates may reflect regional epidemiological characteristics. Second, the automated switching software was developed specifically for the VITEK MS PLUS system; its generalizability requires further validation in diverse laboratory settings. Furthermore, technical considerations warrant careful attention. When yeast is inadvertently processed using the bacterial mode without the addition of formic acid, spectrum acquisition may be suboptimal, necessitating cautious interpretation of the results. Additionally, this study did not evaluate the performance of the databases for identifying filamentous fungi, Mycobacteria or Nocardia species, which represent distinct technical challenges in MALDI-TOF MS analysis (11, 31). Finally, while rare, misidentifications did occur with the RUO database. Therefore, alongside stringent quality control, the clinical interpretation of novel or exceedingly rare species identified by the RUO database must be correlated with epidemiological data and the patient’s clinical presentation.

The “IVD Primary Screening – Automated RUO Supplementation” workflow is designed to be regulatory-compliant, prioritizing approved IVD results while conditionally invoking the RUO database as a supplementary tool. This hierarchical, trigger-based approach enables laboratories to safely maximize technological resources within the current regulatory framework (32). The integrated protocol achieved a stable overall identification rate of 98.7%, indicating that most pathogens can be identified rapidly on-site, reducing delays associated with external referral. This supports timely management of critical infections such as sepsis (33). Looking ahead, our model points toward a more personalized and intelligent diagnostic pathway. As databases expand and AI algorithms advance, future systems may automatically select optimal databases and assess the clinical relevance of identification results. We recommend that laboratories adopting similar RUO databases perform local performance verification, establish concordance benchmarks and develop clear SOPs for reporting and review. Such efforts also contribute to the broader one health framework that integrates human, animal and environmental health.

## Conclusion

The VITEK MS IVD database delivers standardized, high-throughput identification for routine pathogens, while the RUO database provides essential supplementary coverage for challenging isolates and enables workflow optimization, forming a complementary diagnostic toolkit. We recommend that clinical laboratories implement an integrated “IVD primary screening first, RUO targeted supplementation” strategy supported by automated switching software. This approach maintains accuracy and compliance while maximizing identification coverage, speed and cost-effectiveness, ultimately enhancing precise infectious disease management.

## Acknowledgements

Sequencing service was provided by Personal Biotechnology Co., Ltd. Shanghai, China.

## Author contributions

HN, FL, XW contributed equally to this work. HN: Supervision, Formal analysis, Writing – original draft, Visualization, Data curation, Validation, Conceptualization, Writing review & editing. YC: Visualization, Validation, Conceptualization, Writing – review & editing, Writing – original draft, Data curation. FL: Data curation, Visualization, Conceptualization, Validation, Writing – review & editing, Writing – original draft. JD: Visualization, Validation, Writing – review & editing, Writing – original draft. XW: Conceptualization, Validation, Writing – review & editing, Writing – original draft. XY: Data curation, Visualization, Conceptualization, Validation, Writing – original draft.

## Ethics statement

Collection of samples was approved by the First hospital of China Medical University of Ethics Committee for Medical Science Research (No.【2025】151). Isolates from human clinical specimens were enrolled under a waiver of consent.

## Data availability statement

The raw sequencing data have been published under the GenBank, and accession numbers are provided in Table S1

## References

1 Calderaro A, and Chezzi C. 2024. MALDI-TOF MS: A Reliable Tool in the Real Life of the Clinical Microbiology Laboratory. Microorganisms 12 10.3390/microorganisms12020322

2 Chen XF, Hou X, Xiao M, Zhang L, Cheng JW, Zhou ML, Huang JJ, Zhang JJ, Xu YC, and Hsueh PR. 2021. Matrix-Assisted Laser Desorption/Ionization Time of Flight Mass Spectrometry (MALDI-TOF MS) Analysis for the Identification of Pathogenic Microorganisms: A Review. Microorganisms 9 10.3390/microorganisms9071536

3 Kostrzewa M, Nagy E, Schrottner P, and Pranada AB. 2019. How MALDI-TOF mass spectrometry can aid the diagnosis of hard-to-identify pathogenic bacteria - the rare and the unknown. Expert Rev Mol Diagn 19:667–682. 10.1080/14737159.2019.1643238

4 Lee H, Koo J, Oh J, Cho SI, Lee H, Lee HJ, Sung GH, and Kim J. 2024. Clinical Evaluation of VITEK MS PRIME with PICKME Pen for Bacteria and Yeasts, and RUO Database for Filamentous Fungi. Microorganisms 12 10.3390/microorganisms12050964

5 Hou T-Y, Chiang-Ni C, and Teng S-H. 2019. Current status of MALDI-TOF mass spectrometry in clinical microbiology. Journal of food and drug analysis 27:404–414.

6 Tsuchida S, and Nakayama T. 2022. MALDI-based mass spectrometry in clinical testing: focus on bacterial identification. Applied Sciences 12:2814.

7 Daley CL, Flume PA, van Ingen J, Khare R, Mingora CM, Nguyen MH, Winthrop KL, and Zha BS. 2025. A systematic review of tetracyclines for nontuberculous mycobacteria: focus on rapidly growing mycobacteria. Eur Respir Rev 34 10.1183/16000617.0038-2025

8 Alobaid K, Asadzadeh M, Bafna R, and Ahmad S. 2021. First Isolation of Candida nivariensis, an Emerging Fungal Pathogen, in Kuwait. Med Princ Pract 30:80–84. 10.1159/000511553

9 Li Y, Gan Z, Zhou X, and Chen Z. 2022. Accurate classification of Listeria species by MALDI-TOF mass spectrometry incorporating denoising autoencoder and machine learning. J Microbiol Methods 192:106378. 10.1016/j.mimet.2021.106378

10 Church D, Griener T, and Gregson D. 2025. Multi-year comparison of VITEK MS performance for identification of rarely encountered pathogenic gram-positive organisms (GPOs) in a large integrated Canadian healthcare region. Microbiol Spectr 13:e0254524. 10.1128/spectrum.02545-24

11 Rodriguez -Temporal D, Zvezdanova ME, Benedi P, Marin M, Blazquez-Sanchez M, Ruiz-Serrano MJ, Munoz P, and Rodriguez-Sanchez B. 2023. Identification of Nocardia and non-tuberculous Mycobacterium species by MALDI-TOF MS using the VITEK MS coupled to IVD and RUO databases. Microb Biotechnol 16:778–783. 10.1111/1751-7915.14146

12 Topic Popovic N, Kazazic SP, Bojanic K, Strunjak-Perovic I, and Coz-Rakovac R. 2023. Sample preparation and culture condition effects on MALDI-TOF MS identification of bacteria: A review. Mass Spectrom Rev 42:1589–1603. 10.1002/mas.21739

13 Martiny D, Busson L, Wybo I, El Haj RA, Dediste A, and Vandenberg O. 2012. Comparison of the Microflex LT and Vitek MS systems for routine identification of bacteria by matrix-assisted laser desorption ionization-time of flight mass spectrometry. J Clin Microbiol 50:1313–1325. 10.1128/JCM.05971-11

14 Tsuchida S, Umemura H, and Nakayama T. 2020. Current Status of Matrix-Assisted Laser Desorption/Ionization-Time-of-Flight Mass Spectrometry (MALDI-TOF MS) in Clinical Diagnostic Microbiology. Molecules 25 10.3390/molecules25204775

15 Leyer C, Gregorowicz G, Mougari F, Raskine L, Cambau E, and de Briel D. 2017. Comparison of Saramis 4.12 and IVD 3.0 Vitek MS Matrix-Assisted Laser Desorption Ionization-Time of Flight Mass Spectrometry for Identification of Mycobacteria from Solid and Liquid Culture Media. J Clin Microbiol 55:2045–2054. 10.1128/JCM.00006-17

16 Grenfell RC, da Silva Junior AR, Del Negro GM, Munhoz RB, Gimenes VM, Assis DM, Rockstroh AC, Motta AL, Rossi F, Juliano L, Benard G, and de Almeida Junior JN. 2016. Identification of Candida haemulonii Complex Species: Use of ClinProTools(TM) to Overcome Limitations of the Bruker Biotyper(TM), VITEK MS(TM) IVD, and VITEK MS(TM) RUO Databases. Front Microbiol 7:940. 10.3389/fmicb.2016.00940

17 Bachli P, Baars S, Simmler A, Zbinden R, and Schulthess B. 2022. Impact of MALDI-TOF MS identification on anaerobic species and genus diversity in routine diagnostics. Anaerobe 75:102554. 10.1016/j.anaerobe.2022.102554

18 Wayne P. 2017. Clinical and Laboratory Standards Institute (CLSI) Methods for the identification of cultured microorganisms using matrix-assisted laser desorption/ionization time-offlight mass spectrometry. CLSI guideline M58.

19 Wang CH, Su YS, and Lee WS. 2023. Cyberlindnera fabianii fungemia complicating psoas muscle abscess successfully treated by surgical drainage and echinocandin therapy. J Microbiol Immunol Infect 56:644–646. 10.1016/j.jmii.2022.11.001

20 Al-Sweih N, Ahmad S, Khan S, Joseph L, Asadzadeh M, and Khan Z. 2019. Cyberlindnera fabianii fungaemia outbreak in preterm neonates in Kuwait and literature review. Mycoses 62:51–61. 10.1111/myc.12846

21 Turan D, Habip Z, Odabasi H, Dombekci E, Gundogus N, Ozmen M, and Aksaray S. 2025. Antifungal Susceptibilities of Rare Yeast Isolates. J Fungi (Basel) 11 10.3390/jof11090645

22 Malek M, Mrowiec P, Klesiewicz K, Skiba-Kurek I, Szczepanski A, Bialecka J, Zak I, Bogusz B, Kedzierska J, Budak A, and Karczewska E. 2019. Prevalence of human pathogens of the clade Nakaseomyces in a culture collection-the first report on Candida bracarensis in Poland. Folia Microbiol (Praha) 64:307–312. 10.1007/s12223-018-0655-7

23 Merlino J, Rizzo S, Beresford R, and Gray T. 2023. Isolation of Acinetobacter bereziniae harbouring plasmid bla(NDM-1) in central Sydney, Australia. Pathology 55:867–868. 10.1016/j.pathol.2023.02.008

24 Martinez V, Matabang MA, Miller D, Aggarwal R, and LaFortune A. 2023. First case report on Empedobacter falsenii bacteremia. IDCases 33:e01814. 10.1016/j.idcr.2023.e01814

25 Panigrahi MK, Kaliaperumal V, Akella A, Venugopal G, and Ramadass B. 2022. Mapping microbiome-redox spectrum and evaluating Microbial-Redox Index in chronic gastritis. Sci Rep 12:8450. 10.1038/s41598-022-12431-x

26 Obermayer A, Davis J, Talada DP, Teng M, Eschrich S, Yin V, Spakowicz D, Chatterjee D, Rounbehler RJ, Churchman ML, Tarhini AA, Wang X, Gupta S, Markowitz J, Goecks J, Li R, Rodrigues Pessoa R, Manley BJ, Tan AC, Grass GD, Chen DT, and Shaw TI. 2026. ShinyEvents: harmonizing longitudinal data for real-world survival estimation. NPJ Precis Oncol 10.1038/s41698-025-01212-0

27 Jan HE, Lo CL, Lee JC, Li MC, Lin WL, Ko WC, and Lee NY. 2023. Clinical impact of the combination of rapid species identification and antifungal stewardship intervention in adults with candidemia. J Microbiol Immunol Infect 56:1253–1260. 10.1016/j.jmii.2023.08.014

28 Jeon YD, Seong H, Kim D, Ahn MY, Jung IY, Jeong SJ, Choi JY, Song YG, Yong D, Lee K, Kim JM, and Ku NS. 2018. Impact of matrix-assisted laser desorption/ionization time of flight mass spectrometric evaluation on the clinical outcomes of patients with bacteremia and fungemia in clinical settings lacking an antimicrobial stewardship program: a pre-post quasi experimental study. BMC Infect Dis 18:385. 10.1186/s12879-018-3299-y

29 Dunnam G, Thornton JK, and Pulido-Landinez M. 2023. Characterization of an Emerging Enterococcus cecorum Outbreak Causing Severe Systemic Disease with Concurrent Leg Problems in a Broiler Integrator in the Southern United States. Avian Dis 67:137–144. 10.1637/aviandiseases-D-22-00085

30 Li N, Dou S, Feng L, Zhu Q, and Lu N. 2020. Eliminating sweet spot in MALDI-MS with hydrophobic ordered structure as target for quantifying biomolecules. Talanta 218:121172. 10.1016/j.talanta.2020.121172

31 Pastrone L, Curtoni A, Criscione G, Scaiola F, Bottino P, Guarrasi L, Iannaccone M, Timke M, Costa C, and Cavallo R. 2023. Evaluation of Two Different Preparation Protocols for MALDI-TOF MS Nontuberculous Mycobacteria Identification from Liquid and Solid Media. Microorganisms 11 10.3390/microorganisms11010120

32 Mortier T, Wieme AD, Vandamme P, and Waegeman W. 2021. Bacterial species identification using MALDI-TOF mass spectrometry and machine learning techniques: A large-scale benchmarking study. Comput Struct Biotechnol J 19:6157–6168. 10.1016/j.csbj.2021.11.004

33 Xu X, Wang Z, Lu E, Lin T, Du H, Li Z, and Ma J. 2025. Rapid detection of carbapenem-resistant Escherichia coli and carbapenem-resistant Klebsiella pneumoniae in positive blood cultures via MALDI-TOF MS and tree-based machine learning models. BMC Microbiol 25:44. 10.1186/s12866-025-03755-5

